# Spatio-temporal regulation of endocytic protein assembly by SH3 domains in yeast

**DOI:** 10.1101/2022.08.11.503583

**Authors:** Daniel R. Hummel, Marko Kaksonen

## Abstract

Clathrin-mediated endocytosis is a conserved eukaryotic membrane trafficking pathway that is driven by a sequentially assembled molecular machinery that contains over 60 different proteins. SH3 domains are the most abundant protein-protein interaction domain in this process, but the function of most SH3 domains in protein dynamics remains elusive. Using mutagenesis and live-cell fluorescence microscopy in the budding yeast *Saccharomyces cerevisiae*, we dissected SH3-mediated regulation of the endocytic pathway. Our data suggest that multiple SH3 domains regulate the actin nucleation promoting Las17/Vrp1 complex, and that the network of SH3 interactions coordinate both Las17/Vrp1 assembly and dissociation. Furthermore, most endocytic SH3 domain proteins use the SH3 domain for their own recruitment, while a minority uses the SH3 domain to recruit other proteins, and not themselves. Our results provide a dynamic map of SH3 functions in yeast endocytosis and a framework for SH3 interaction network studies across biology.

## Introduction

Clathrin-mediated endocytosis is a conserved eukaryotic process by which cells internalize plasma membrane and extracellular material. Over 60 different proteins assemble at the plasma membrane to coordinate cargo collection, formation of a membrane invagination and vesicle release (McMahon & Boucrot, 2011; Kaksonen & Roux, 2018). In the budding yeast *Saccharomyces cerevisiae*, endocytic protein assembly is stereotypical and sequential. The endocytic proteins can be categorized based on their spatio-temporal dynamics at endocytic sites into one of four modules (Kaksonen et al., 2005): the coat, the actin nucleation promoting factors (NPF), the actin patch or the scission module. However, within a module, individual proteins characteristically assemble and disassemble with different kinetics, which implies that the protein-protein interactions (PPI) between endocytic proteins occur within a precise spatial and temporal context. This symphony of events is poorly understood, but central to endocytic site progression. A major PPI network in endocytosis is the SH3 interaction network (Tong et al., 2002; Tonikian et al., 2009). SH3 domains are a versatile PPI domain family, typically found in eukaryotic proteins involved in pathways regulating the cytoskeleton and membrane traffic. In yeast endocytosis, SH3 domains are the most abundant PPI domain family and the only domain family present in all four endocytic modules (Goode et al., 2015). The two major categories of functions of endocytic SH3 interactions are the coordination of actin regulators and protein recruitment (Fazi et al., 2002; Tong et al., 2002; Sun et al., 2006; Tonikian et al., 2009; Feliciano & Di Pietro, 2012; Sun et al., 2015; Sun et al., 2017).

Yeast endocytosis is dependent on the formation of a branched actin network, which drives membrane invagination (Kaksonen et al., 2003). The primary actin nucleation promoting factors (NPFs), found in the NPF module, are the WASp-homolog, Las17, and the paralogous type-I myosins Myo3 and Myo5 in conjunction with verprolin, Vrp1 (Weinberg & Drubin, 2012; Goode et al., 2015). Las17 is in a complex with Vrp1 and together this complex contains numerous proline-rich motifs, which are considered to be binding targets of the SH3 domains of all nine endocytic SH3 proteins in different modules: Sla1, Lsb3/4 paralogs (coat), Bzz1, Bbc1, Myo3/5 paralogs (NPF), Abp1 (actin) and Rvs167 (scission) (Tong et al., 2002; Tonikian et al., 2009). Sla1 recruits Las17 to endocytic sites via its SH3 domains (Feliciano & Di Pietro, 2012; Sun et al., 2015). Las17 molecules localize to a ring-shaped region around the coat (Mund et al., 2018). During the actin phase, Sla1 is associated with the invagination tip, but the Las17-Vrp1 complex (LVC) is located at the invagination base where it is regulated by Bbc1 and Myo5 (Kaksonen et al., 2005; Sun et al., 2006; Picco et al., 2015). Bbc1 limits the Las17 ring size, and inhibits an excessive assembly of actin filaments (Mund et al., 2018; Picco et al., 2018), possibly via SH3 interactions. The Myo5 SH3 domain binds Vrp1, which promotes Myo5 assembly and thereby actin network growth (Anderson et al., 1998; Sun et al., 2006). These data point to a central role for SH3 domains regulating actin network assembly at endocytic sites. Evidence also indicates that the SH3 domains of Myo5, Lsb4 and Rvs167 are involved in the recruitment of these proteins to endocytic sites (Sun et al., 2006; Urbanek et al., 2015; Menon & Kaksonen, 2021). This is in contrast to Sla1, which has an SH3-independent recruitment to endocytic sites (Sun et al., 2015). Whether the other SH3 proteins use their SH3 domains to regulate the LVC complex or their own recruitment remains an open question.

In this study, we aimed to connect SH3-mediated SH3 protein assembly with SH3-mediated LVC regulations. We tested if assembly of Lsb3, Bzz1, Bbc1 and Abp1 is dependent on their SH3 domains. Our results suggest a new categorization of endocytic SH3 domains, which is based on their functions as recruitment factors. Furthermore, we show that Abp1, Bzz1 and Rvs167 SH3 interactions regulate the LVC at endocytic sites. These new LVC regulations may indicate feedback mechanisms for actin nucleation control.

## Results

### The SH3 domains of Bzz1 have distinct functions

Bzz1 is an NPF module protein with two SH3 domains of unknown function. We aimed to dissect if and how these SH3 domains regulate Bzz1 assembly. From the N-to C-terminus Bzz1 has a membrane-binding F-BAR domain, an SH3 domain (SH3A), a proline-rich motif (PRM) and a second SH3 domain (SH3B) (fig. 1A). Metazoan orthologs, Pascins/Syndapins and Fchsds/Nervous wreck, have reported intramolecular interactions between their F-BAR and SH3 domains (Rao et al., 2010; Stanishneva-Konovalova et al., 2016). A similar membrane specificity and SH3-mediated PPI may also regulate Bzz1 assembly. A previous report suggests that F-BAR membrane binding promotes Bzz1 recruitment but a role for the Bzz1 SH3 domains is undescribed (Kishimoto et al., 2011).

**Figure 1:**
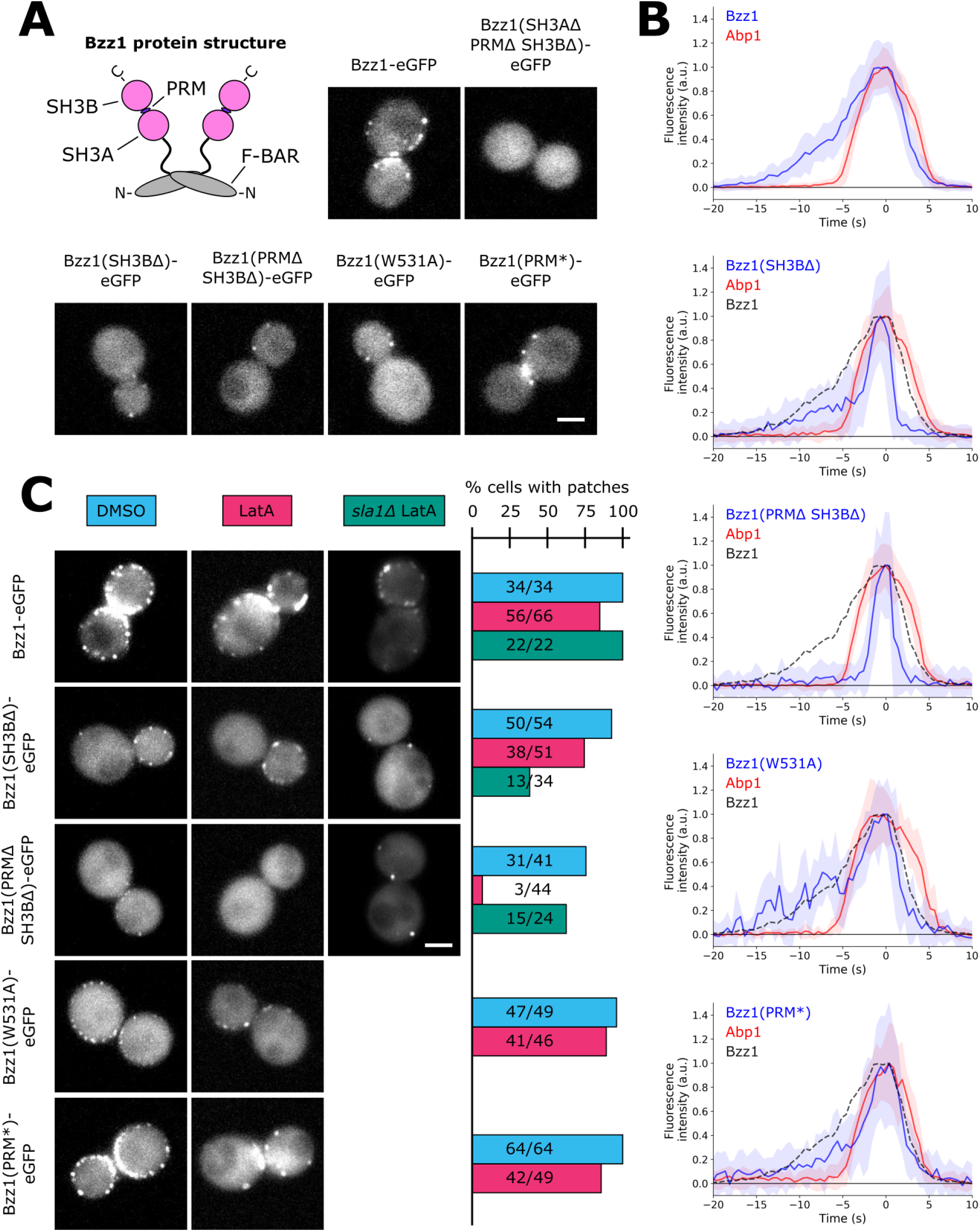
Bzz1’s SH3 domains regulate Bzz1 recruitment distinctly.**A)** Scheme of Bzz1 protein structure and localization of WT and mutant Bzz1 proteins. Scale bar: 2 μm. **B)** Average fluorescence signals of WT and mutant Bzz1 and Abp1. Solid lines are medians, shaded areas are median absolute deviation scaled for asymptotically normal consistency. Number of patches used to generate the average: Bzz1: 68, Bzz1(SH3BΔ): 53, Bzz1(PRMΔ SH3BΔ): 38, Bzz1(SH3A*): 13 and Bzz1(PRM*): 26. **C)** Bzz1 patch formation upon LatA treatment in WT and *sla1Δ* cells. Images of representative cells are maximum intensity projections of 5 min time lapse movies. Scale bar: 2 μm. Quantification shows the percentage of cells with patches.

Bzz1 SH3 domains were investigated by endogenously-tagged Bzz1 mutant proteins with a C-terminal eGFP and imaged by live-cell fluorescence microscopy. Bzz1-eGFP formed patches at the plasma membrane (fig. 1A) similar to previous reports (Sun et al., 2006). In contrast, Bzz1 protein lacking both SH3 domains and the PRM (Bzz1(SH3AΔ PRMΔ SH3BΔ)-eGFP) showed no patch formation at all (fig. 1A), which suggests that its C-terminal domains are needed for Bzz1 recruitment to endocytic sites. Next, we tested the contribution of individual C-terminal domains to Bzz1 recruitment. We analyzed two truncation mutants, Bzz1(SH3BΔ)-eGFP and Bzz1(PRMΔ SH3BΔ)-eGFP, a point mutant, Bzz1(W531A)-eGFP, which altered the canonical binding site of the Bzz1 SH3A domain, and a PRM mutant Bzz1(PRM*)-eGFP, where the PRM was substituted with a flexible linker (PAPEVPPPRR → SGGGGSGGGG). Each of these four Bzz1 mutants formed patches (fig. 1A). To be more sensitive to low intensity fluorescence, we used total internal reflection fluorescence (TIRF) microscopy to quantify Bzz1 assembly dynamics. In addition, we used Abp1-mCherry as an actin marker and time reference for the endocytic actin phase. Bzz1-eGFP had a lifetime of about 20 seconds (fig. 1B), comparable to previous reports (Sun et al., 2006; Kishimoto et al., 2011). Abp1-mCherry started assembling about 10 s after Bzz1, reached the peak intensity together with Bzz1 and then the two proteins disassembled with similar timing. Bzz1 assembly rate was constant or slightly exponential until Bzz1 molecule numbers reached their maximum. In contrast, before the Abp1 signal, Bzz1(SH3BΔ)-eGFP assembled slowly but then during the actin phase accumulated more quickly. Bzz1(PRMΔ SH3BΔ)-eGFP only assembled during the actin phase and its signal resembled the second, fast accumulation phase of Bzz1(SH3BΔ). Bzz1(W531A)-eGFP signal was similar to the Bzz1-eGFP signal. The assembly slope of Bzz1(PRM*)-eGFP was similar to the one of Bzz1(SH3BΔ), with a slow start and a quick accumulation during the actin phase. The data suggest that the PRM and SH3B interactions regulate initial Bzz1 assembly before the actin phase and the SH3A promotes Bzz1 recruitment specifically during the actin phase.

We suspected that the Bzz1 SH3A interaction could be actin-dependent due to its specific contribution to Bzz1 assembly during the actin phase. We tested if the Bzz1 mutants form patches in cells treated with Latrunculin A (LatA), which inhibits actin polymerization and consequently invagination growth, effectively stalling endocytic sites (Ayscough et al., 1997; Kaksonen et al., 2003). Loss of Abp1-mCherry patches was used as a control for the efficacy of the LatA treatment. Upon LatA treatment, all Bzz1 mutants retained patch localization except Bzz1(PRMΔ SH3BΔ)-eGFP (fig. 1C). Thus, the Bzz1 SH3A interaction requires the actin phase to occur, while the PRM and SH3B interactions are actin-independent.

Previous studies have suggested that Sla1 SH3 domains compete with Bzz1 SH3 domains (Sun et al., 2006) and others indicate Sla1 SH3 interactions with the Bzz1 PRM (Barry & Di Pietro, 2015). Consequently, actin-dependency of Bzz1 patch formation could change in the absence of Sla1. Hence, we treated *sla1Δ* cells with LatA and analyzed WT and mutant Bzz1 patch formation. We noted a change in the percentage of cells that retained Bzz1 patches and we quantified this phenotype. Bzz1-eGFP patch occurrence was slightly increased by the deletion of Sla1 (fig. 1C). In the case of Bzz1(SH3BΔ)-eGFP, patch formation could be observed in only around half as many LatA-treated *sla1Δ* than WT cells (fig. 1C). Surprisingly, Bzz1(PRMΔ SH3BΔ)-eGFP, the Bzz1 mutant that did not form patches in LatA-treated WT cells, formed patches in the majority of LatA-treated *sla1Δ* cells (fig. 1C). Thus, Bzz1’s SH3A can promote Bzz1 recruitment in LatA-treated cells in the absence of Sla1. This observation can also explain the patch formation of Bzz1(SH3BΔ)-eGFP in LatA-treated *sla1Δ* cells. Our results suggest that the Bzz1 PRM interacts with Sla1 SH3 domains but also that Sla1 inhibits Bzz1’s SH3A. In conclusion, to promote Bzz1 assembly, the Bzz1 SH3A interaction requires the actin phase in the presence of Sla1, but not in absence of Sla1. These results support the binding competition between Sla1 and Bzz1 for Las17 observed *in vitro* and suggest these competition dynamics change during the actin phase.

### Bzz1 and Rvs167 antagonize Sla1 and promote Las17 dissociation

Deletion of Bzz1 causes only mild endocytic phenotypes (Sun et al., 2006; Kishimoto et al., 2011), which complicates the analysis of its function. We speculated that Bzz1’s deletion phenotype may be masked by other redundant regulatory pathways. The observation that Bzz1 is inhibited by Sla1 might therefore be a gateway to revealing Bzz1’s function in endocytosis. We speculated that Bzz1 might reveal a function in LatA-treated *sla1Δ* cells as the Bzz1 SH3 interactions are not inhibited in this condition. Thus, we analyzed patch formation of potential Bzz1 interaction partners in LatA-treated WT and *sla1Δ* cells. The NPF module proteins Las17, Vrp1 and Myo5 formed persistent patches in LatA-treated WT cells when observed over a period of 5 minutes (fig. 2A). However, in LatA-treated *sla1Δ* cells, patch formation of Las17, Vrp1 and Myo5 was disturbed (fig. 2A). We could observe three types of patches for the NPF module proteins: (A) dim and persistent patches at the plasma membrane, (B) bright and transient patches at the plasma membrane and (C) transient patches in the cytoplasm. Las17, Vrp1 and Myo5 appeared to be unstably associated with endocytic sites in LatA-treated *sla1Δ* cells. In contrast, the coat protein, Pan1, only formed persistent patches at the plasma membrane in both LatA-treated WT and *sla1Δ* cells (fig. 2A). These data suggest that specifically NPF module dissociation is misregulated in LatA-treated *sla1Δ* cells. We hypothesized that the non-inhibited Bzz1 causes the dissociation of the NPF module from endocytic sites. Thus, we analyzed Las17 patch formation in LatA-treated WT, *sla1Δ* and *sla1Δ bzz1Δ* cells (fig. 2B):

**Figure 2:**
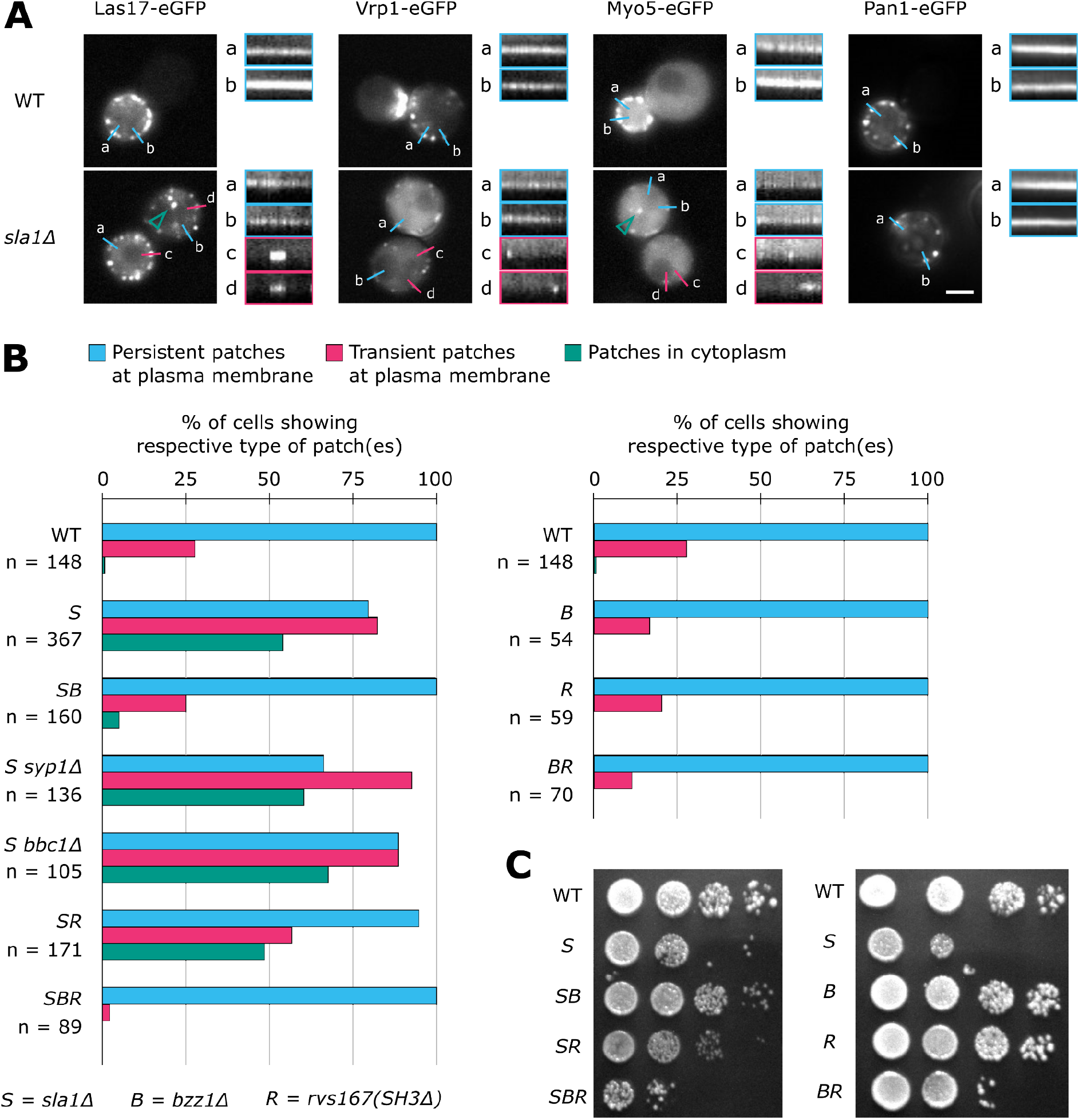
Bzz1 and Rvs167 regulate NPF association with the coat in absence of Sla1.**A)** LatA treatment of WT and *sla1Δ* cells. Example kymographs of persistent patches (a) and b)) and of transient patches (c) and d)). Pictures of representative cells are maximum intensity projections of 4 minutes and 50 seconds long time lapses. Scale bar: 2 μm. Kymograph length: 4 min 50 seconds. **B)** Bzz1 and Rvs167 perturb Las17 association to the plasma membrane in a Sla1-dependent fashion. In LatA treated *sla1Δ* cells, Las17-eGFP forms persistent (blue) and transient (pink) patches at the plasma membrane as well as cytoplasmic patches (green). Data from 2 to 6 experiments per strain was pooled together. The total number of cells analyzed is indicated by n. The time lapses analyzed had a duration of 4 minutes 50 seconds and a frame rate of 10 seconds. **C)** Spot growth assay. Cells were grown 48 hours at 30 °C. All cells express Las17-eGFP and Abp1-mCherry endogenously. Abbreviations: *S* = *sla1Δ, B* = *bzz1Δ* and *R* = *rvs167(SH3Δ)*.

The number of cells showing persistent Las17 patches decreased from 100 % in LatA-treated WT cells to 80 % in LatA-treated *sla1Δ* cells, but was again 100 % in *sla1Δ bzz1Δ* cells. Transient Las17 patch formation at the plasma membrane could be observed in 28 % of LatA-treated WT cells. For LatA-treated *sla1Δ* cells, it was 82 %, roughly three times more compared to WT cells. However, in LatA-treated *sla1Δ bzz1Δ* cells, the number of cells with transient plasma membrane-associated patches was 25 %, close to WT levels. Cytoplasmic Las17 patches could be observed in just 1 % of LatA-treated WT cells but in 54 % of the LatA-treated *sla1Δ* cells. In contrast to that, only 5 % of the LatA-treated *sla1Δ bzz1Δ* cells showed cytoplasmic Las17 patches. These results suggested that, in absence of Sla1, Bzz1 decreases the formation of stable Las17 patches, promotes transient Las17 patch formation and affects Las17 localization. Overall, deletion of Bzz1 largely rescued normal Las17 patch association to the plasma membrane in LatA-treated *sla1Δ* cells. Taken together these data indicate that Bzz1 perturbs Las17 association to endocytic sites and that this function can be inhibited by Sla1.

We then tested whether deletion of other proteins could also rescue the Las17 phenotype observed in LatA-treated *sla1Δ* cells. We chose Syp1 and Bbc1, two other patch proteins that regulate Las17 (Weinberg & Drubin, 2012; Goode et al., 2015), as well as Rvs167 due to its domain similarity to Bzz1. Deletion of Rvs167 causes failure of endocytic vesicle scission and cell growth defects. Therefore, we used a deletion of the Rvs167 SH3 domain, *rvs167(SH3Δ)*, which causes a milder phenotype (Menon & Kaksonen, 2021).

In LatA-treated *sla1Δ syp1Δ* and *sla1Δ bbc1Δ* cells, the number of cells with either transient plasma membrane-associated or cytoplasmic patches were comparable to *sla1Δ* cells (fig. 2B). In contrast, the deletion of the Rvs167 SH3 domain partly suppressed the Las17 phenotype, but the effect was milder than the deletion of Bzz1 (fig. 2B). These results suggest that Rvs167 and Bzz1 both regulate Las17 in absence of Sla1. Therefore we combined the Bzz1 and Rvs167 mutations and analyzed Las17 patches in LatA-treated *sla1Δ bzz1Δ rvs167(SH3Δ)* cells. We found that Las17 patches were hyper-persistent as the percentage of cells with transient Las17 patches at the plasma membrane was only 2 % in LatA-treated *sla1Δ bzz1Δ rvs167(SH3Δ)* cells, much lower than in LatA-treated WT cells (fig. 2B). These results suggest that in *sla1Δ* cells Bzz1 and Rvs167 can modify Las17 patch persistence. When Sla1 was not deleted, the effect of *bzz1Δ, rvs167(SH3Δ)* and *bzz1Δ rvs167(SH3Δ)* on the number of cells with transient Las17 patches at the plasma membrane was weak (fig. 2B). Thus, it appears that Sla1 largely inhibits the effects of Bzz1 and Rvs167 in regard to Las17 regulation in LatA-treated cells.

To see if Bzz1 and Rvs167 antagonize Sla1 also in normal drug free conditions, we tested whether growth defects of *sla1Δ* cells could be restored by the deletion of Bzz1 and/or the Rvs167 SH3 domain (fig. 2C). The *sla1Δ bzz1Δ* and *sla1Δ rvs167(SH3Δ)* strains indeed grew much better than *sla1Δ* cells, which confirmed that Bzz1 and Rvs167 antagonism to Sla1 is not an artifact of the LatA-treatment. The deletion of Bzz1 had a slightly stronger effect on growth rescue than the deletion of the Rvs167 SH3 domain, correlating with the effects on Las17 patch formation in LatA-treated cells (fig. 2B). The combined mutant *sla1Δ bzz1Δ rvs167(SH3Δ)* grew slightly worse in comparison with *sla1Δ* cells. The single deletion of Bzz1 or the Rvs167 SH3 domain showed no effect on cell growth, but *bzz1Δ rvs167(SH3Δ)* cells had a synthetic growth defect (fig. 2C). Cooperation of Bzz1 and Rvs167 has already been suggested, but in the context of vesicle scission (Kishimoto et al., 2011). Our growth assay suggests a commonality between Bzz1 and Rvs167 in addition to scission control, as *rvs167(SH3Δ)* causes no scission failure (Menon & Kaksonen, 2021).

In summary, our results show that Las17 association to endocytic sites is disturbed in LatA-treated *sla1Δ* cells. Deletion of the Rvs167 SH3 domain and Bzz1 could rescue this defect and also partially restore growth defect of *sla1Δ* cells. Thus, our results suggest that Bzz1 and Rvs167 have an antagonistic function to Sla1 and can act as Las17 disassembly factors.

### Las17/Vrp1 positioning along the invagination axis is regulated by the Abp1 SH3 domain

The antagonistic relationship between Sla1 and Bzz1/Rvs167 suggests that different SH3 domain interactions with Las17 can modify each other. Abp1, the SH3 protein of the actin module, assembles before Rvs167 and the Bzz1 SH3A interaction. An indication for potential SH3 crosstalk is the synthetic lethality of Abp1 and Sla1 mutants (Holtzman et al., 1993; Haynes et al., 2007). Abp1 has an N-terminal actin-binding domain and a C-terminal SH3 domain. First, we tested whether the Abp1 SH3 domain mediates Abp1 assembly. Abp1(SH3Δ)-eGFP formed apparently normal plasma membrane -associated patches like Abp1-eGFP, but also motile patches that were not associated with the plasma membrane (fig. 3A). Furthermore, we could observe bigger Abp1(SH3Δ)-eGFP structures that looked like clusters of patches. Hence, deletion of the Abp1 SH3 domain affected Abp1 localization.

**Figure 3:**
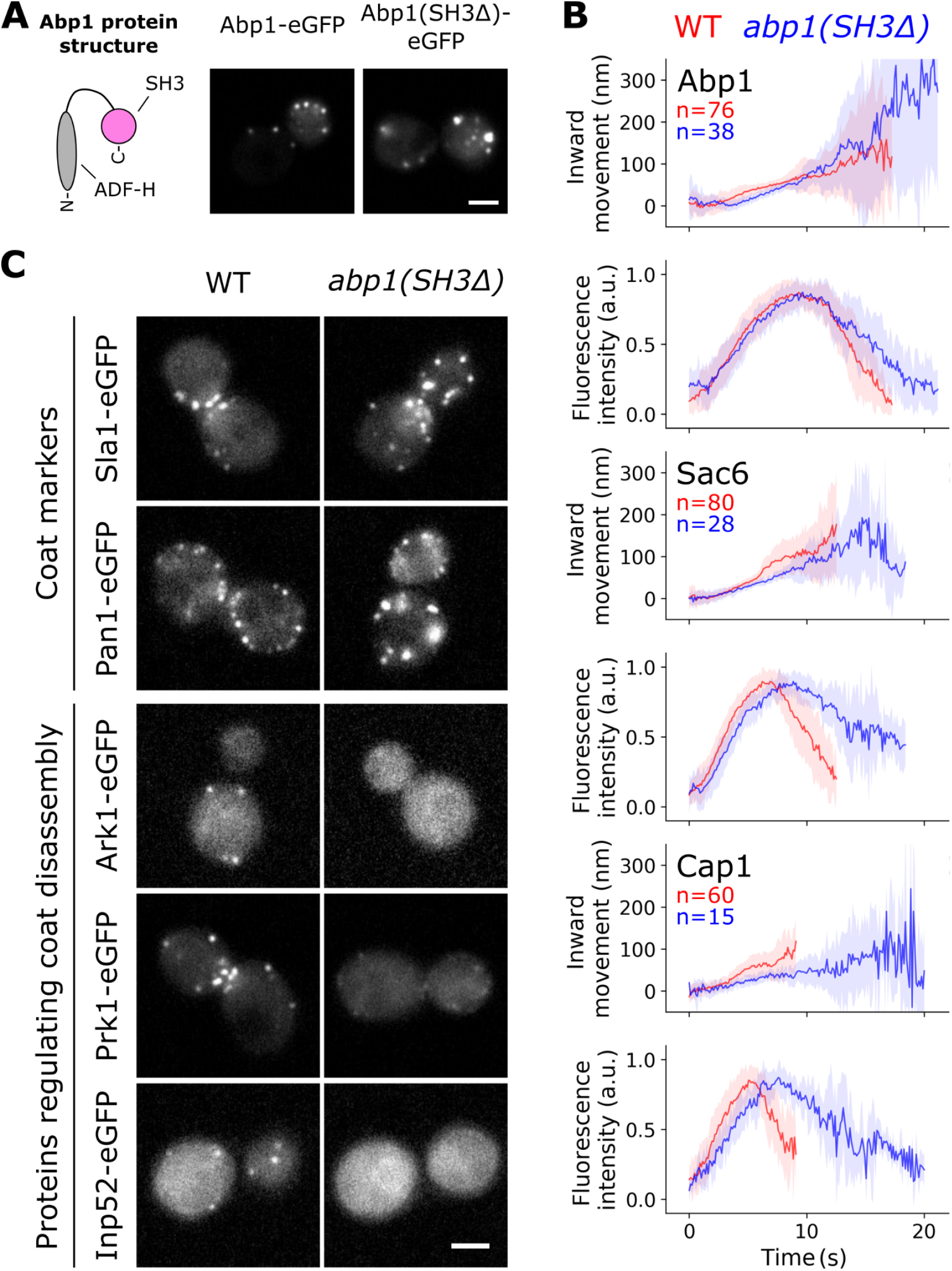
The Abp1 SH3 domain recruits coat disassembly factors and mediates actin disassembly dynamics.**A)** Abp1(SH3Δ) assembles at endocytic sites but also forms cytoplasmic clusters. **B)** Median patch movement and lifetimes of Abp1 and Abp1(SH3Δ) as well as Sac6 and Cap1 in WT (red) and *abp1(SH3Δ)* (blue) cells. Depicted are median centroid movement and fluorescence intensities (solid lines) with median absolute deviations (shaded areas). **C)** Top: Coat proteins form cytosolic clusters in *abp1(SH3Δ)* cells. Bottom: Deletion of the Abp1 SH3 domain inhibits recruitment of the coat disassembly regulators Ark1, Prk1 and Inp52. Pictures of representative cells are single frames. Scale bar: 2 μm

We analyzed the centroid movement and lifetimes of Abp1-eGFP and Abp1(SH3Δ)-eGFP patches that were associated with the plasma membrane (fig. 3B). The deletion of its SH3 domain did not affect Abp1 assembly, but slowed down Abp1 disassembly dynamics relative to full length Abp1. We could observe a similar phenotype for the actin module proteins, Sac6 and Cap1, in *abp1(SH3Δ)* cells (fig. 3B). Thus, it is likely that the Abp1 SH3 domain does not regulate Abp1 disassembly specifically, but generally affects actin network disassembly dynamics.

Similar to Abp1(SH3Δ), the coat proteins, Sla1 and Pan1, also formed cytoplasmic patches and clusters in *abp1(SH3Δ)* cells (fig. 3C). That is reminiscent of disassembly defects reported for Ark1, Prk1 and Inp52 mutant cells (Cope et al., 1999; Sekiya-Kawasaki et al., 2003; Stefan et al., 2005; Toret et al., 2008). Deletion of the Abp1 SH3 domain strongly inhibited Ark1, Prk1 and Inp52 patch formation (fig. 3C), in agreement with previous reports (Fazi et al., 2002; Stefan et al., 2005). In Ark1, Prk1 and Inp52 mutant cells many endocytic proteins fail to disassemble from the newly made vesicle and thereby inhibit further trafficking steps (Toret et al., 2008). The cytoplasmic patches and clusters of Abp1, Sla1 and Pan1 in *abp1(SH3Δ)* cells may reflect a similar vesicle trafficking blockage.

The Abp1 SH3 interactions therefore regulate the endocytic machinery post-vesicle scission, in particular for vesicle uncoating and actin disassembly. To investigate potential Abp1 SH3 functions prior to scission, we analyzed Rvs167-eGFP patches in WT and *abp1(SH3Δ)* cells. Rvs167 is a scission marker and altered Rvs167 assembly dynamics reflect defects in invagination growth and/or scission (Kukulski et al., 2012; Picco et al., 2015). In WT cells, Rvs167 assembled for around 5 seconds, then molecule numbers rapidly dropped and simultaneously the patch centroid moved quickly inward (fig. 4A), which correlates with vesicle scission. Rvs167 patch movement in *abp1(SH3Δ)* cells suggests that the final invagination size was similar to WT cells (fig. 4A). However, vesicle scission was delayed and occurred 3 seconds later in the mutants. Furthermore, like coat and actin module proteins (fig. 4B & C), Rvs167 also formed aberrant cytoplasmic structures in *abp1(SH3Δ)* cells.

**Figure 4:**
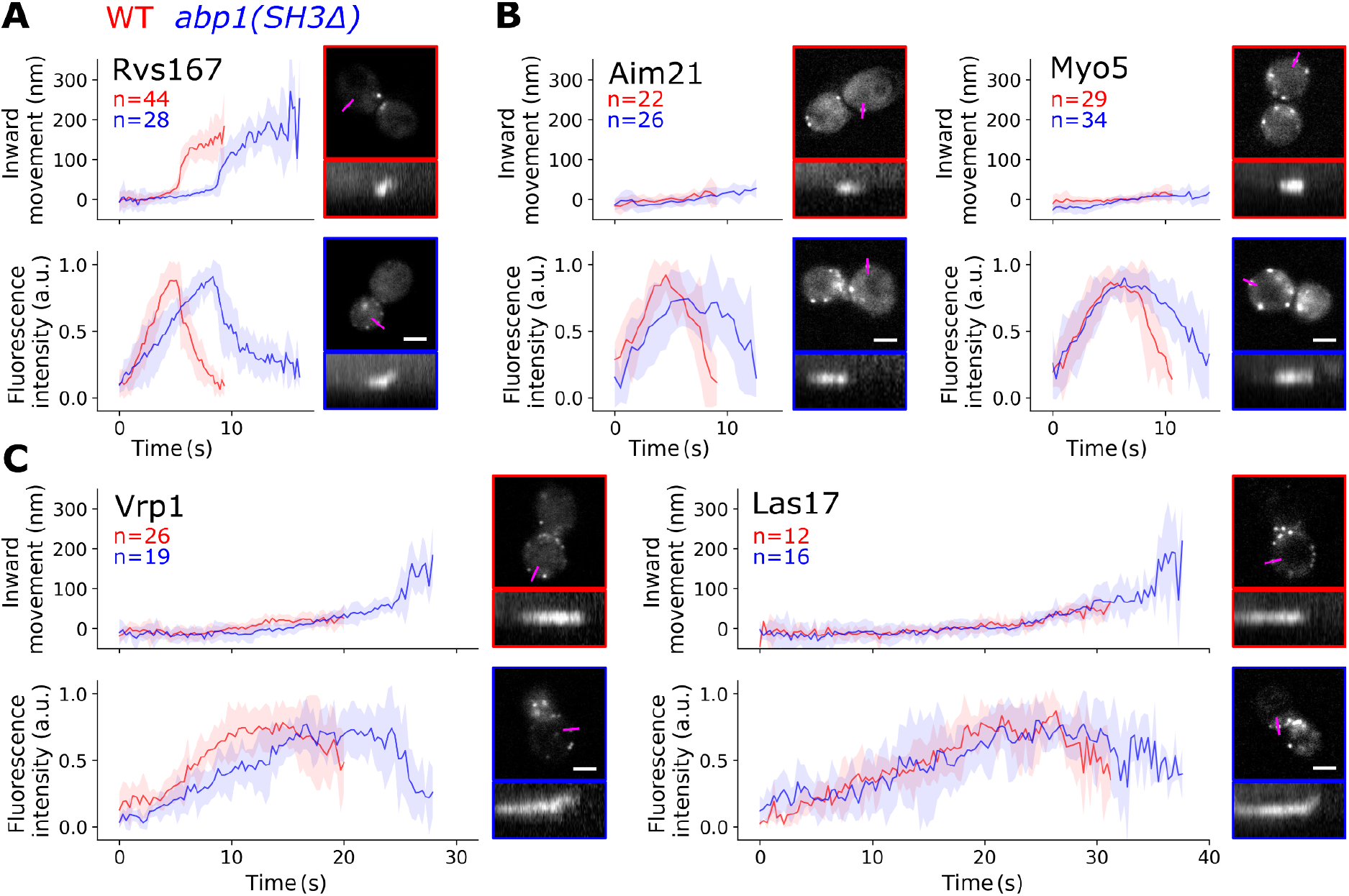
Abp1 SH3 domain interactions promote scission, NPF disassembly and LVC anchoring.**A)** Patch movement and lifetimes of Rvs167 in WT and *abp1(SH3Δ)* cells. **B)** Patch movement and lifetimes of Aim21 and Myo5 in WT and *abp1(SH3Δ)* cells. **C)** Patch movement and lifetimes of Vrp1 and Las17 in WT and *abp1(SH3Δ)* cells. Plots show median centroid movement and fluorescence intensities (solid lines) with median absolute deviations (shaded areas). Pictures of representative cells are single frames. Scale bar: 2 μm. Kymograph length: 36 seconds.

We then analyzed patch dynamics of the NPF module proteins Las17, Vrp1, Myo5 and Aim21 in WT and *abp1(SH3Δ)* cells. All four proteins formed cytosolic patches, similar to the actin patch and coat proteins (fig. 4B & C), suggesting that NPFs might promote actin nucleation at cytosolic patches and thereby enhance actin disassembly defects even further. Regarding endocytic patches at the plasma membrane, Myo5 and Aim21 patch lifetimes were around 3 to 4 seconds longer in the mutant than in WT cells (fig. 4B), similar to the delay in vesicle scission. Also Vrp1 and Las17 lifetimes were longer in *abp1(SH3Δ)* than in WT cells (fig. 4C). On average, Vrp1 had a lifetime of around 20 seconds and Las17 of around 30 seconds in WT cells. In *abp1(SH3Δ)* cells, lifetimes were about 8 seconds longer for both Vrp1 and Las17. Furthermore, Las17 and Vrp1 patches formed non-motile patches in WT cells. Strikingly, however, in *abp1(SH3Δ)* cells, membrane-associated Las17 and Vrp1 patches were motile at the end of their lifetimes (fig. 4C). This indicates a defect in Las17 and Vrp1 association to the invagination base and to the myosins which form non-motile patches in both WT and *abp1(SH3Δ)* cells (fig. 4B). De-localization of the LVC from the myosins effectively splits the actin nucleation promoting complex.

Association of proteins to blocked vesicles might reduce the pool of proteins available for endocytosis, affecting protein assembly dynamics in *abp1(SH3Δ)* cells. This could explain the extended lifetimes of Rvs167, Aim21, Myo5, Vrp1 and Las17 at endocytic sites in these cells. However, the mechanism causing Las17 and Vrp1 patch movement in *abp1(SH3Δ)* cells remains unclear. The Abp1 SH3 domain could directly or indirectly modify upstream regulators of Las17 and Vrp1 like Sla1, mediate simultaneous interaction partners like Myo5, or promote downstream interactions with Bzz1 and Rvs167.

### Lsb3 and Bbc1 SH3 domain-dependent recruitment

Our data shows that Bzz1 SH3 domains regulate Bzz1 assembly (fig. 1). Recruitment of Rvs167 and Myo5 is also mediated by their SH3 domains, as reported previously (Sun et al., 2006; Menon & Kaksonen, 2021). The Myo3 SH3 domain likely acts similarly to the paralogous Myo5, recruiting Myo3. In contrast, Sla1 (Sun et al., 2015) and Abp1 (fig. 3A & B) do not appear to use their SH3 domains for their own recruitment to endocytic sites. We aimed to characterize the remaining SH3 domains of Lsb3/4 and Bbc1 in regard to protein recruitment.

We focused on Lsb3 as a proxy for the Lsb3/4 paralogs due to issues with endogenous tagging of Lsb4 (Dewar et al., 2002; Urbanek et al., 2015). Deletion of the Lsb3 SH3 domain reduced Lsb3 patch intensities strongly but did not prevent patch formation entirely (fig. 5A). In LatA-treated cells, both Lsb3 and Lsb3(SH3Δ) formed patches (fig. 5A). Hence, Lsb3 recruitment seems to be actin-independent with two recruitment routes, one via its N-terminal domain and one via the SH3 interaction.

**Figure 5:**
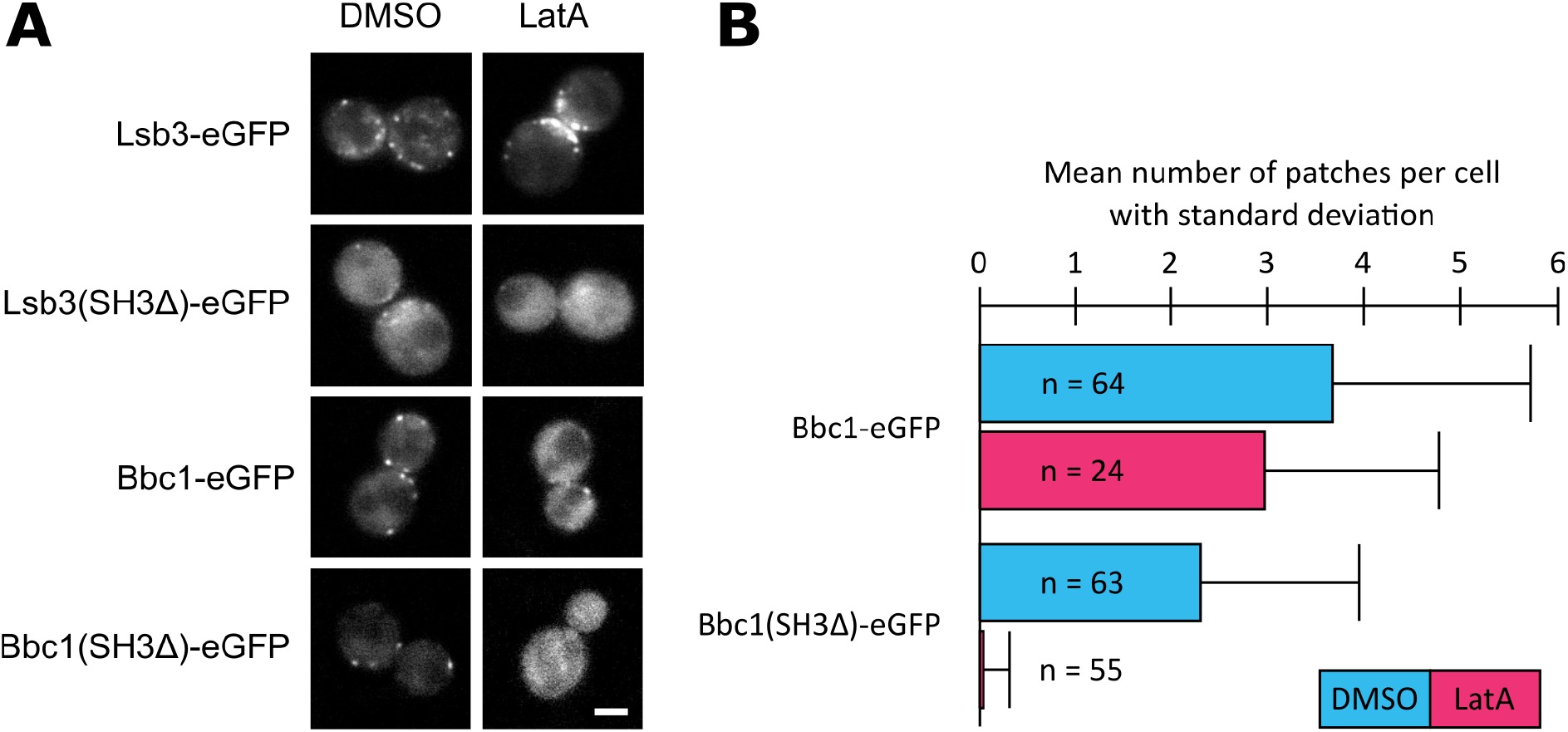
The Lsb3 and Bbc1 SH3 domain regulate Lsb3 and Bbc1 recruitment, respectively.**A)** Patch formation of Lsb3, Lsb3(SH3Δ), Bbc1 and Bbc1(SH3Δ) in LatA treated cells. Scale bar: 2 μm. **B)** Mean number of Bbc1 and Bbc1(SH3Δ) patches per cell.

In the case of Bbc1, the deletion of its SH3 domain also reduced protein patch intensities and increased diffuse cytosolic signals (fig. 5A). Furthermore, we could observe a reduction in the number of patches per cell for Bbc1(SH3Δ) compared to Bbc1 (fig. 5B). Upon LatA treatment, Bbc1 formed patches but Bbc1(SH3Δ) did not (fig. 5A & B). Thus, the Bbc1 SH3 domain promotes Bbc1 recruitment in an actin-independent fashion while the rest of Bbc1 has actin-dependent interactions.

Taken together, Lsb3 and Bbc1 each have SH3 domains that regulate their own protein recruitment to endocytic sites, similarly to the SH3 domains of Myo5, Rvs167 and Bzz1.

## Discussion

### Spatio-temporal regulation of the LVC by the SH3 network

Sla1, Myo5 and Bbc1 are established regulators of the LVC (Rodal et al., 2003; Kaksonen et al., 2005; Sun et al., 2006; Feliciano & Di Pietro, 2012; Weinberg & Drubin, 2012; Mund et al., 2018; Picco et al., 2018). We found that the SH3 domain proteins Bzz1, Rvs167 and Abp1 are also regulators of the LVC at endocytic sites, emphasizing the central nature of the dynamic SH3 PPI network. The complexity of the SH3 PPI network has hampered the understanding of its function in regulating endocytosis. We propose three different perspectives on the SH3 domain-mediated dynamic regulation of the LVC. Together with previous studies our results suggest roles for SH3 domains in temporal coordination (fig. 6A), spatial coordination (fig. 6B) and a dynamic network coordination of LVC regulation (fig. 6C) at endocytic sites.

**Figure 6:**
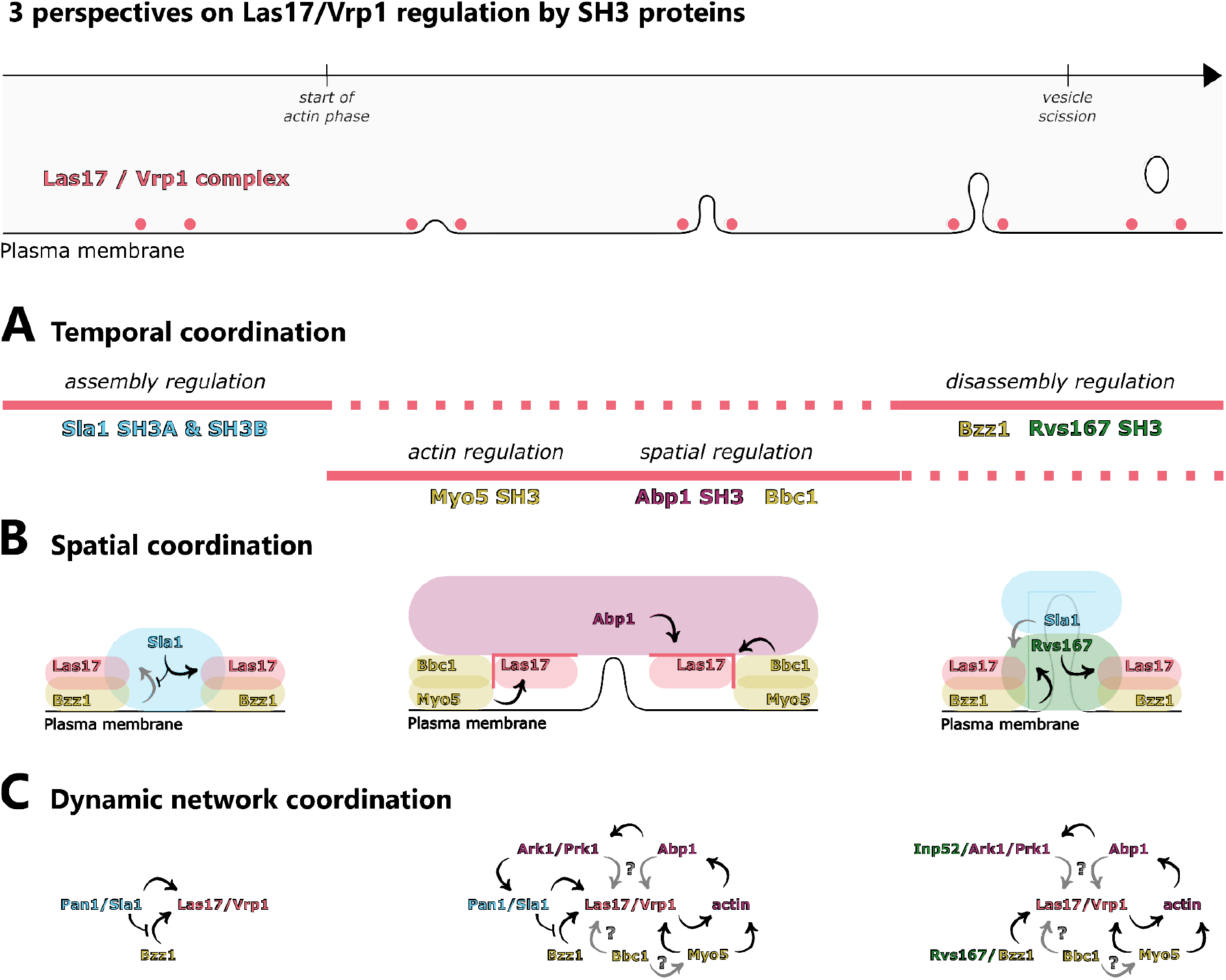
Conclusions and models about LVC regulations by SH3 domains in yeast endocytosis. Top: LVC lifetime and localization during a stereotypical endocytic event.**A**) SH3 proteins regulate the LVC assembly dynamics and localization over time. **B)** Relative localization of SH3 proteins within the endocytic machinery regulates the LVC and its SH3 interactions. For readability, only key features of the respective phase are depicted. **C)** Dynamics of the endocytic SH3 interaction network. Color code: Salmon: LVC, blue = coat, yellow = NPF module, pink = actin module, green = scission module.

The LVC is initially recruited to endocytic sites before actin assembly occurs. The initial assembly of the LVC is regulated by Sla1. Sla1 has three SH3 domains (SH3A, SH3B and SH3C). The Sla1 SH3A and SH3B recruit Las17 by binding a central part of the Las17 proline-rich region (Feliciano & Di Pietro, 2012; Sun et al., 2015). During actin assembly, the Myo5 SH3 domain interacts with Vrp1 and Myo5 regulates actin together with the LVC (Sun et al., 2006). Simultaneously, Bbc1 inhibits excessive recruitment of Las17 to the periphery of the NPF ring structure (Mund et al., 2018). Concurrent with these activities, the Abp1 SH3 domain regulates LVC localization along the invagination axis (fig. 4). As Bbc1 and Abp1 mediate LVC positioning within the endocytic machinery, we consider them “spatial regulators” of the LVC. Eventually, at the final stage of the actin phase, formation of new actin filaments slows down just prior to vesicle scission and correlates with LVC disassembly (Picco et al., 2018). Our data indicates that Bzz1 and the Rvs167 SH3 domains coordinate LVC disassembly from endocytic sites (fig. 2).. Spatial regulation of the LVC might support the disassembly regulation by Bzz1 and Rvs167. Furthermore, there may be a smooth change from assembly to disassembly regulation, in particular as Sla1 and Bzz1 appear to be in a binding competition (fig. 1). In conclusion, the LVC undergoes different phases of regulation. These regulatory phases coordinate LVC recruitment, positioning of the LVC within the endocytic machinery and finally disassembly the LVC.

SH3 domains have an intrinsic specificity and different SH3 domains preferentially bind different targets (Tong et al., 2002; Tonikian et al., 2009). However, swapping of SH3 domains, even within one protein, influences those SH3 domains’ specificity (Dionne et al., 2021). Hence, molecular context defines SH3 interactions, too, and the molecular context is constantly changing during endocytic site progression. In a flat membrane scenario, Sla1 inhibits Bzz1 SH3 interactions that promote Bzz1 assembly during the later actin phase (fig. 1). Interactions of Bzz1’s F-BAR domain with the plasma membrane may affect Bzz1 SH3 competition with Sla1, perhaps in a membrane composition- or curvature-dependent fashion (Takano et al., 2008; Rao et al., 2010; Stanishneva-Konovalova et al., 2016). In addition to membrane curvature, invagination growth exposes Bzz1 to a number of proteins that have changed their positions relative to each other. We suspect that these architectural changes of the endocytic machinery during invagination growth affect SH3 interactions (fig. 6B). During invagination growth, the LVC and Bzz1 are located at the invagination base but Sla1 is associated with the invagination tip (Weinberg & Drubin, 2012). This physical distancing could perturb Sla1 interactions and consequently change Sla1-Bzz1 competition dynamics. In addition, Rvs167 assembles at the invagination neck close to the LVC, which is where Sla1 is located until the start of membrane invagination (Picco et al., 2015). Rvs167 may interfere with Sla1 interactions and cooperate with Bzz1 to regulate LVC disassembly (fig.2). The interplay of membrane curvature and relative changes in protein positions change endocytic sites from favoring Sla1 SH3 domain interactions to favoring Bzz1 and Rvs167 SH3 domain interactions. At the same time, Myo5, Bbc1 and Abp1 regulate the LVC, too. While Sla1, Bzz1 and Rvs167 are located on the inside of the LVC ring, Myo5, Bbc1 and Abp1 assemble outside of it (Mund et al., 2018). The spatial regulators Bbc1 and Abp1 are positioned like a barrier around the LVC (fig. 6B), potentially regulating SH3 interactions that depend on LVC ring geometry. The LVC moves from its normal position in *abp1(SH3Δ)* cells (fig. 4), maybe because it remains bound to Sla1. The displacement of the LVC could allow it to escape from Bzz1- and Rvs167-mediated disassembly, explaining slow LVC disassembly in *abp1(SH3Δ)* cells. In conclusion, molecular context seems to affect SH3 interactions with the LVC and might provide feedback mechanisms for LVC regulation.

Temporal (fig. 6A) and spatial (fig. 6B) aspects of LVC regulation by SH3 proteins imply that the SH3 interactions change dynamically during the progression of endocytosis (Figure 6C). Initially, Sla1 interacts with the LVC. At this stage the Bzz1 PRM and SH3B promote Bzz1 recruitment. Sla1 competes with the Bzz1 SH3A, effectively inhibiting its interactions with the LVC (fig. 1). With the start of the actin phase, however, the SH3 interactions network becomes bigger and more complex. Myo5 and Bbc1 regulate the LVC but also interact with each other (Mochida et al., 2002). The LVC and the myosins mediate the formation of a branched actin network, which recruits Abp1. The Abp1 SH3 domain recruits Ark1 and Prk1 (fig. 3) (Fazi et al., 2002), leading to disassembly of the Sla1/Pan1 complex. Effectively, the circle closes and the network becomes a self-regulating process based on negative feedback. The Ark1/Prk1-mediated disassembly of the Sla1/Pan1 complex in combination with Sla1 moving away from the LVC, probably reduces the importance of Sla1 interactions when Rvs167 and Inp52 interactions occur (Menon & Kaksonen, 2021). The coat disassembly factor Inp52 would down-regulate any residual Sla1 interaction even further. Due to the reduced Sla1-mediated inhibition, Bzz1 and Rvs167 could promote LVC disassembly. Subsequently, actin nucleation ends and LVC-dependent proteins dissociate from endocytic sites. This dynamic SH3 interaction network appears to self-regulate by feedback mechanisms. Such a tangled system may be important to allow adaptation to changes in cellular physiology or environmental conditions and could be a major reason for the remarkable robustness of endocytosis.

### SH3 domains as recruitment factors - recruiting or recruited?

We can now compare the endocytic SH3 domains in regard to their function as recruitment factors. As depicted in table 1, Sla1 and Abp1 do not use their SH3 domains for their own recruitment. However, these two proteins are known to recruit other endocytic components via their SH3 domains (Fazi et al., 2002; Stefan et al., 2005; Feliciano & Di Pietro, 2012; Sun et al., 2015). In contrast, Lsb3/4, Bzz1, Bbc1, Myo3/5 and Rvs167 use their SH3 domains to regulate their own recruitment (fig. 1 & 5)(Sun et al., 2006; Urbanek et al., 2015; Menon & Kaksonen, 2021). For this group of SH3 proteins there is so far no evidence that they would recruit other proteins using their SH3 domains. As SH3 domains typically have only one binding site, it is likely that one SH3 domain can fulfill only one recruitment function at a time. Thus, the SH3 domains are either recruiting their own protein or another protein, but not both. This type of recruitment specificity might be a useful generalization for understanding the functions of SH3 domains in PPI networks, such as the human one with hundreds of SH3 proteins.

**Table 1:** SH3 domains as recruitment factors.

|  |  | SH3 domain recruits other proteins? |  |
| --- | --- | --- | --- |
|  |  | Yes | Absence of evidence |
| SH3 domain regulates assembly of its own protein? | Yes |  | Lsb3/4, Bzz1, Bbc1, Myo3/5, Rvs167 |
|  | No | Sla1, Abp1 |  |

## Author contributions

D.R. Hummel and M. Kaksonen designed the experiments. D.R. Hummel performed the experiments and analyzed the data. D.R. Hummel and M. Kaksonen wrote the manuscript. M. Kaksonen secured the funding.

## Acknowledgments

We are thankful to all the members of the Kaksonen laboratory, especially Andrea Picco for advice on data analysis and Python scripting, and Christopher Toret for critical reading of the manuscript. Furthermore, we would like to thank Florian Shala and Loris Levet for technical help. This work was supported by the Swiss National Science Foundation (grant 310030B_182825) and by the NCCR Chemical Biology funded by the SNSF.

## Methods

### Strains, media and plasmids

Strains were created via homologous recombination with PCR cassettes, and by mating and sporulation. EGFP-tagging, deletions and truncations were confirmed by PCR, substitutions by sequencing. All fluorescently tagged genes are expressed endogenously.

### Live cell imaging

#### Sample preparation

Yeast cells were grown to OD600 between 0.3 and 0.8 at 24 ºC with shaking in Synthetic Complete medium without L-Tryptophan (SC-TRP). 40 μl of cells were added to a coverslip coated with Concanavalin A. After 5 to 10 minutes of incubation, cells were washed with fresh SC-TRP medium, 40 μl of fresh SC-TRP medium was added and the cells were transferred to the microscope. For DMSO or Latrunculin A (LatA) treatment, cells were instead washed with SC-TRP medium containing 0.2 mM DMSO or LatA and 40 μl of DMSO or LatA medium was added. After 15 minutes of incubation, cells were washed again with DMSO or LatA medium, 40 μl of DMSO or LatA medium was added and the cells were transferred to the microscope.

#### TIRF microscopy and analysis

TIRF microscopy images were acquired using the Olympus IX83 wide-field microscope equipped with a 150x 1.45 oil objective and an ImageEM X2 EM-CCD camera (Hamamatsu) controlled by the VisiView software (Visitron Systems). For illumination of GFP- and mCherry-tagged proteins, a 488 nm and 561 nm laser were used and the laser angles were controlled by the iLas2 system (Roper Scientific). The emission light went through a dichroic Di03-R488/561-t1−25 × 36 filter set (Semrock Brightline) and was split using the W-view Gemini system (Hamamatsu).

TIRF microscopy data was background subtracted with a median filter using ImageJ (Schneider et al., 2012). Spatially and temporarily clearly distinguishable endocytic patches were manually selected with the round selection tool in ImageJ and the fluorescence intensity over time was measured in both color channels. These fluorescence profiles included background signals from before and after the endocytic event. The data was analyzed by a custom-made script using Python 3 controlled by the Jupyter Notebook (Van Rossum & Drake, 2009; Kluyver et al., 2016). First, the data was normalized over the integral of the fluorescence intensity based on Simpson’s rule. Second, the signals were time-aligned by cross-correlating the Abp1-mCherry signals. Third, the median fluorescence intensity and median absolute deviation adjusted by a factor for asymptotically normal consistency was calculated. Fourth, the median intensities were normalized between 0 and 1 and errors were accordingly propagated. The maximum was set to 1 and the average background intensity was set to 0. To obtain an accurate estimation of the background signal, the local minima and maxima of the median signals after the endocytic event were averaged. For the identification of these local minima and maxima, the first derivative of the median signals was estimated by convolution with an 11-point Savitzky-Golay filter.

#### Epi-fluorescence microscopy and analysis

Epi-fluorescence microscopy images were acquired using the Olympus wide-field microscope IX81 equipped with a 100× 1.45 objective and an ORCA-ER CCD camera (Hamamatsu). As illumination source an X-CITE 120 PC (EXFO) metal halide lamp was used. The excitation and emission light when imaging EGFP- and mCherry-tagged proteins were filtered through an U-MGFPHQ and U-MRFPHQ filter set (Olympus), respectively.

Images were processed in ImageJ (Schneider et al., 2012). Images were corrected for background fluorescence using the rolling ball algorithm of ImageJ and movies were in addition corrected for photobleaching using the simple ratio algorithm of ImageJ. For particle detection and tracking, the Particle Tracker (Sbalzarini and Koumoutsakos, 2005) of the MosaicSuite was used. Average trajectories were obtained as described previously (Picco et al., 2015). Trajectory plots were generated using Python 3 controlled by the Jupyter Notebook (Van Rossum & Drake, 2009; Kluyver et al., 2016).

